# Correlated Motion-Based Residue Network Analysis Reveals the Distal Thermal Activation in Soybean Lipoxygenase

**DOI:** 10.64898/2025.12.05.692688

**Authors:** Yaoyukun Jiang, Jaden P. Cordova, Judith P. Klinman

**Affiliations:** Department of Chemistry, University of California, Berkeley, California 94720, United States; California Institute for Quantitative Biosciences, University of California, Berkeley, California 94720, United States; Department of Bioengineering, University of California, Berkeley, California 94720, United States; Department of Molecular and Cell Biology, University of California, Berkeley, California 94720, United States

## Abstract

Enzyme catalysis has been shown to depend on distal pathways that channel thermal energy from solvent to the active site. In soybean lipoxygenase (SLO), experiments identified a cone-shaped network connecting loop residue Gln322 to Leu546 but not Leu754 in the active site. Here, microsecond molecular dynamics and a correlated motion-based protocol provide an atomistic analysis of such long-range communication. The developed approach enables systematic screening of communication between active site-specific residues that directly contact bound substrate and surface-exposed residues on the protein-solvent interface. In doing so, it provides a deeper molecular insight into experimentally mapped networks by resolving communication trends across diverse conformational ensembles. The simulations recover the experimentally demonstrated thermal initiation loop and the Leu546-directed cone in SLO, exclude the negative-control Ser596, and explain the preference for Leu546 over Leu754 through shorter, more correlated helical pathways. Mutational analysis further reveals the impact of single-site mutations on the network preference between Leu546 and Leu754. These results unify experiments and computation, corroborating an anisotropic channeling of thermal energy in SLO and establishing a general framework for computing distal intra-protein pathways that may enable the thermal activation of enzyme function.

## INTRODUCTION

Enzymes accelerate chemical reactions not only through active-site chemistry but also by harnessing protein dynamics.^1–3^ Recent experiments indicate that solvent collisions excite defined residue networks that channel thermal energy anisotropically to the active site, controlling and reducing the enthalpic barrier for substrate turnover in relation to comparable solution reactions.^4^ These “distal communication pathways” are specific to the reaction being catalyzed and distinct from classical allosteric regulation.

Soybean lipoxygenase (SLO) has emerged as a benchmark system for such studies.^5,6^ Time and temperature-resolved hydrogen deuterium exchange and Stokes shifts revealed that surface loop residues, including Gln322, initiate communication along a cone-shaped network terminating at Leu546, while a nearby site, Leu754, lies off-network.^7^ Note that structural modeling of the complex of substrate with SLO indicates that Leu546 and Leu754 bracket opposing faces of the reactive C-H bond of substrate (Figure 1).^8^ This experimental distinction—on-network versus off-network—established that catalysis in SLO depends on selective distal communication. What remains unclear is how such communication is encoded at the molecular level, how pathways are selected, and how mutations reshape their efficiency.

**Figure 1.**
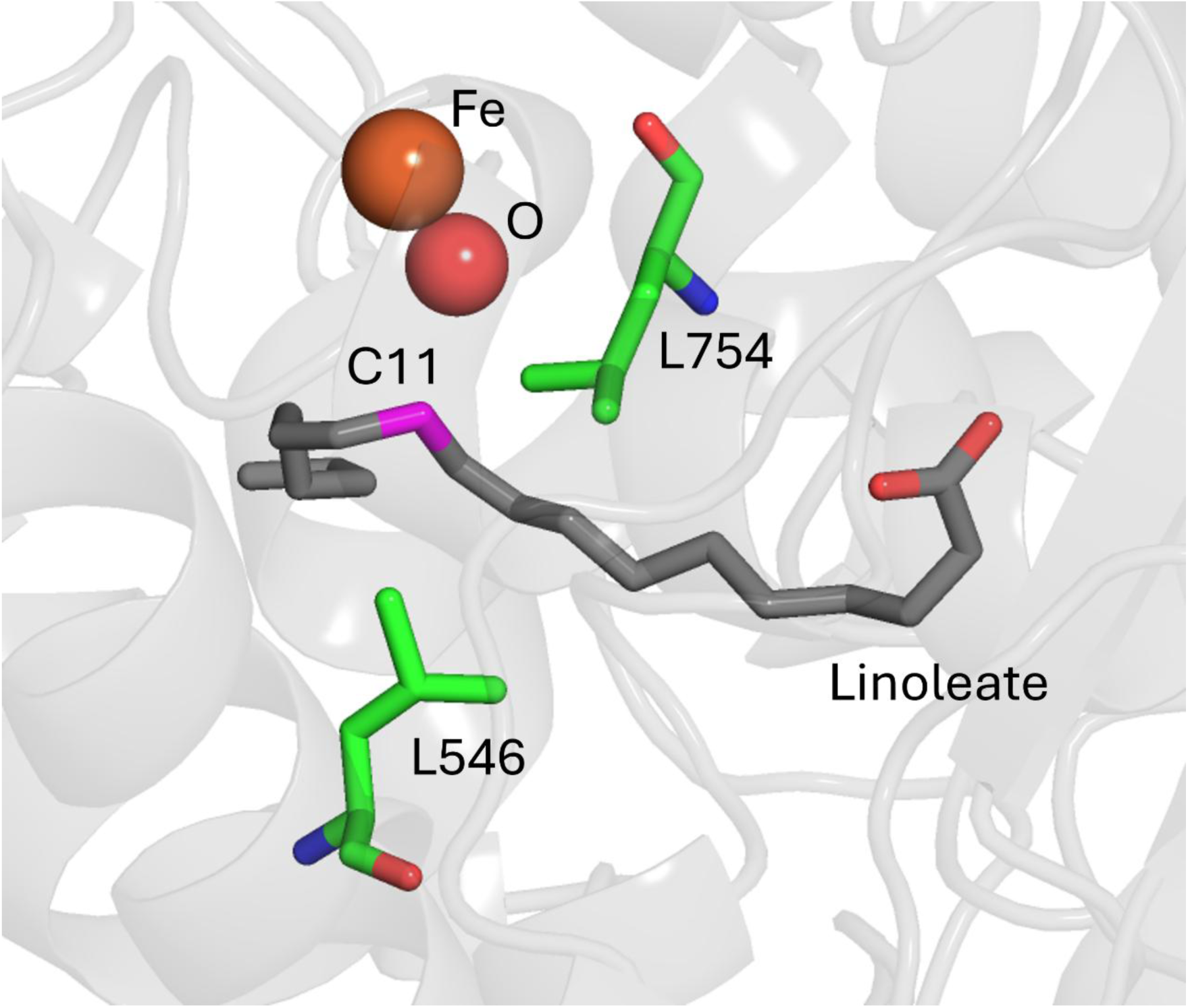
Schematic representation of the active site structure of soybean lipoxygenase. The protein structure is from X-ray crystallography (PDB: 3PZW).^9^ The coordinates of Fe and O are from the crystal structure. The substrate, linoleate, is docked in the apo enzyme from a previous QM/MM study^8^ and is shown in gray sticks. The reactive carbon in the substrate, C11, is highlighted in magenta. The key residues, Leu546 and Leu754, are shown in green sticks. The Fe and bound oxygen are shown as orange and red spheres, respectively.

Here, we address these questions with microsecond molecular dynamics simulations and a correlated motion-based network protocol.^10,11^ This approach resolves residue-level communication pathways between the protein-solvent interface and active site, benchmarks them against the experimentally defined Gln322–Leu546 and alternate Gln322–Leu754 route, and evaluates the impact of site-specific mutations on their relative importance. Our results provide atomistic insight into how SLO preferentially directs energy from the protein-solvent interface to key active site positions, unifying experimental observations with a computational framework for distal intra-enzyme communication.

## RESULTS

### Development of a computational protocol

Molecular dynamics (MD) simulations were performed with the AMBER 24 software suite^12^ using the ff19SB protein^13^ and OPC^14^ water force fields. We simulated SLO without the substrate but with the Fe(II). The starting structure was based on the crystal structure of SLO (PDB ID 3PZW^9^), complemented by another crystal structure (PDB ID 1YGE^15^) for unresolved loops. The Fe(II) center, together with the bound His499, His504, His690, Ile839, and H_2_O, was parameterized with *MCPB.py*.^16^ We assume that treating the active site as Fe(II) bound with H_2_O (as observed from x-ray studies) rather than Fe(III) bound with OH^-^ will have little if any impact on the generic prediction of thermal networks. Proteins were solvated in a truncated octahedron box of ∼32,000 water molecules with sodium counterions. After minimization, systems were heated to 298.15 K, equilibrated under constant pressure, and simulated for 4 μs. Three replicates were conducted for each variant, and snapshots of trajectories were saved every 50 ps.

The developed residue-level network identification protocol to capture communication pathways is summarized in Figure 2. To avoid interdomain interference, only the catalytic domain of SLO (His147–Ile839^17^) was analyzed. Root mean square deviation (RMSD) analysis showed full equilibration after ∼2 μs (Figure S1a). We divided each subsequent 2 μs trajectory into 10 windows, with each window (200 ns) containing 4000 MD frames saved every 50 ps. Individual residues are represented by backbone (BB: C_α_, C, O, N) and side-chain (SC: heavy atoms) nodes, except glycine, which lacks SC. Fe(II) and its bound oxygen are treated as additional nodes, yielding 1349 in total (1348 for I553G).

**Figure 2.**
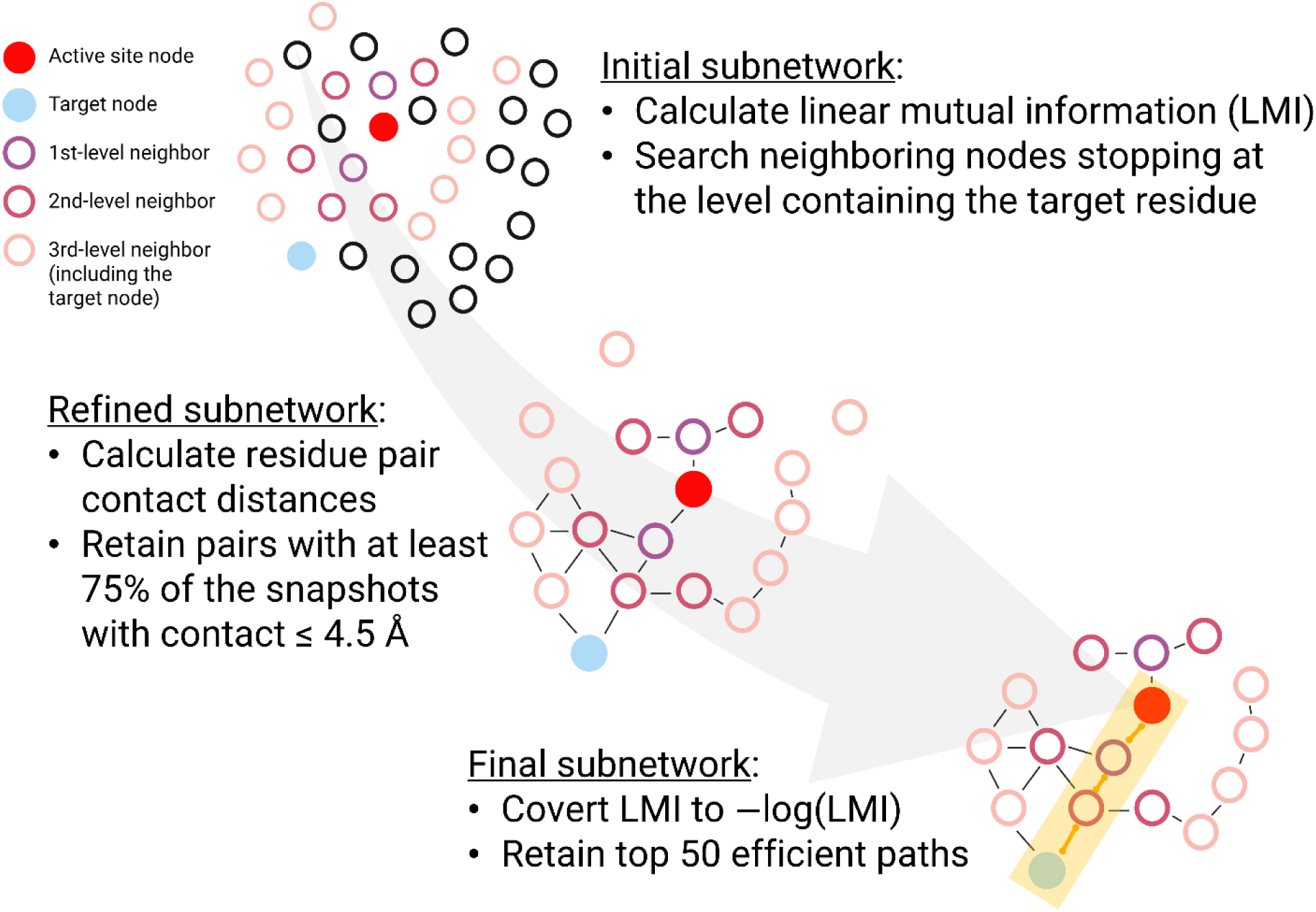
Computational protocol for identifying distal dynamic communication between the protein-solvent interface and the active site, based on networks of correlated motions between persistently contacting residues. The correlated motions are quantified by linear mutual information larger than a dynamically determined cutoff. The initial subnetwork does not involve residue distance evaluation, whereas persistently contacting residues are defined as those with minimum heavy atom distances that are equal to or less than 4.5 Å in at least 75% simulation snapshots. The determination of a network is dependent on an active site side chain adjacent to the bound substrate and a defined residue at the protein-solvent interface. The active site and target nodes are shown in solid red and blue circles, respectively, with intermediate nodes as donuts. For clarity, only three levels of neighboring nodes and one top path (highlighted in orange) are displayed for the final subnetwork.

Correlated motions were first analyzed using linear mutual information (LMI) to quantify the statistical relationship between individual residues,^10,18^ producing a symmetric node–node matrix (Figure S2). It should be noted that LMI only considers the linear portion of the motion correlation. A unity LMI represents perfectly correlated motions, and zero means the absence of correlated motions. Unlike dynamic cross correlation, LMI cannot distinguish correlated (moving in the same direction) from anti-correlated (opposite direction) motions, so the term “correlated motions” used herein includes both cases. To suppress trivial covalent correlations, entries for residues separated by fewer than four sequence positions were removed. Residue connectivity was identified through an iterative neighbor search based on the increasing LMI cutoff. The search herein was initiated with a cutoff of 0.4, and all results between 0.4 and unity were retained. The value for LMI cutoff was then progressively increased until the connectivity between the established source and target residue was lost. The result is subnetworks that link source and target residues with a dynamically determined maximum LMI cutoff.

This subnetwork was refined by proximity-based filtering to ensure persistent physical contacts, where the minimum heavy atom distances are not larger than 4.5 Å in at least 75% of the MD snapshots.^19^ The original quantifier of the edge, LMI, of the refined subnetwork was then converted to a converted LMI (*cLMI*): *cLMI* = − log(*LMI*)^.20^ A lower *cLMI* value represents a higher correlated motion. This value reconciles the goals of finding paths with the highest overall correlated motions and shortest lengths, the latter of which was achieved by the path-finding algorithms (explained in detail in Text S1).^21,22^ The most efficient paths combine high overall correlated motion with the shortest distances. Dijkstra’s algorithm^21^ identified the shortest path between the source and designated nodes. Yen’s algorithm^22^ was then employed to determine the top 49 deviations from this shortest path, if any. This protocol led to at most 200 (50 paths × 4 node combinations) most probable routes within the refined network across all source-terminal backbone/side-chain node combinations (BB-BB, BB-SC, SC-BB, and SC-SC). Note that in practice, we define the source and terminal residues to be in the active site and on the protein-solvent interface, respectively. This is the reverse of a site-specific solvent-derived thermal activation moving toward the active site. Protein-encoded thermal pathways are expected to be intrinsic and independent of the direction analyzed. A detailed description of the simulation and network analysis is provided in Text S1.

### Computationally derived pathways reveal the experimentally observed thermal network of SLO

Using the residue-occurrence analysis described above, we first focused on quantifying communication between the surface Gln322 of the identified loop in SLO and two distal active site residues, Leu546 and Leu754 (Figure 1). The pathway from Gln322 to Leu546 is found to be mediated by fewer residues and shows a higher overall frequency of occurrence compared to Gln322–Leu754 (Figure 3 and Table S1). Specifically, 33,596 events involving 97 residues were observed for Gln322–Leu546, versus 26,686 events across 118 residues for Gln322–Leu754. Applying a cutoff of 600 occurrences (Figure 3b), 14 residues were highly populated for Gln322–Leu546 communication, compared to only 8 residues for Gln322–Leu754. The total occurrence of those 14 residues for Gln322–Leu546 is 19278, compared to only 8098 for those 8 residues for Gln322–Leu754. Focusing only on the endpoints, Gln322 and Leu546 showed 2250 and 2937 occurrences, respectively, compared to 1402 and 1750 for Gln322–Leu754 (Table S1). When residues are restricted to the experimental network previously reported (Leu262, Phe270, Leu299, Ile307, Ile313, Leu316, Tyr317, Gln322, Leu325, Leu546, Ile552, Ile553, Leu742, Ile746, Ser749, and Val750),^7,23^ the cumulative occurrence remained higher for Gln322–Leu546 (6563 vs 4832, excluding Leu546 and Gln322). Together, these results indicate that SLO more efficiently sustains dynamic distal communication between Gln322 and Leu546 than between Gln322 and Leu754.

**Figure 3.**
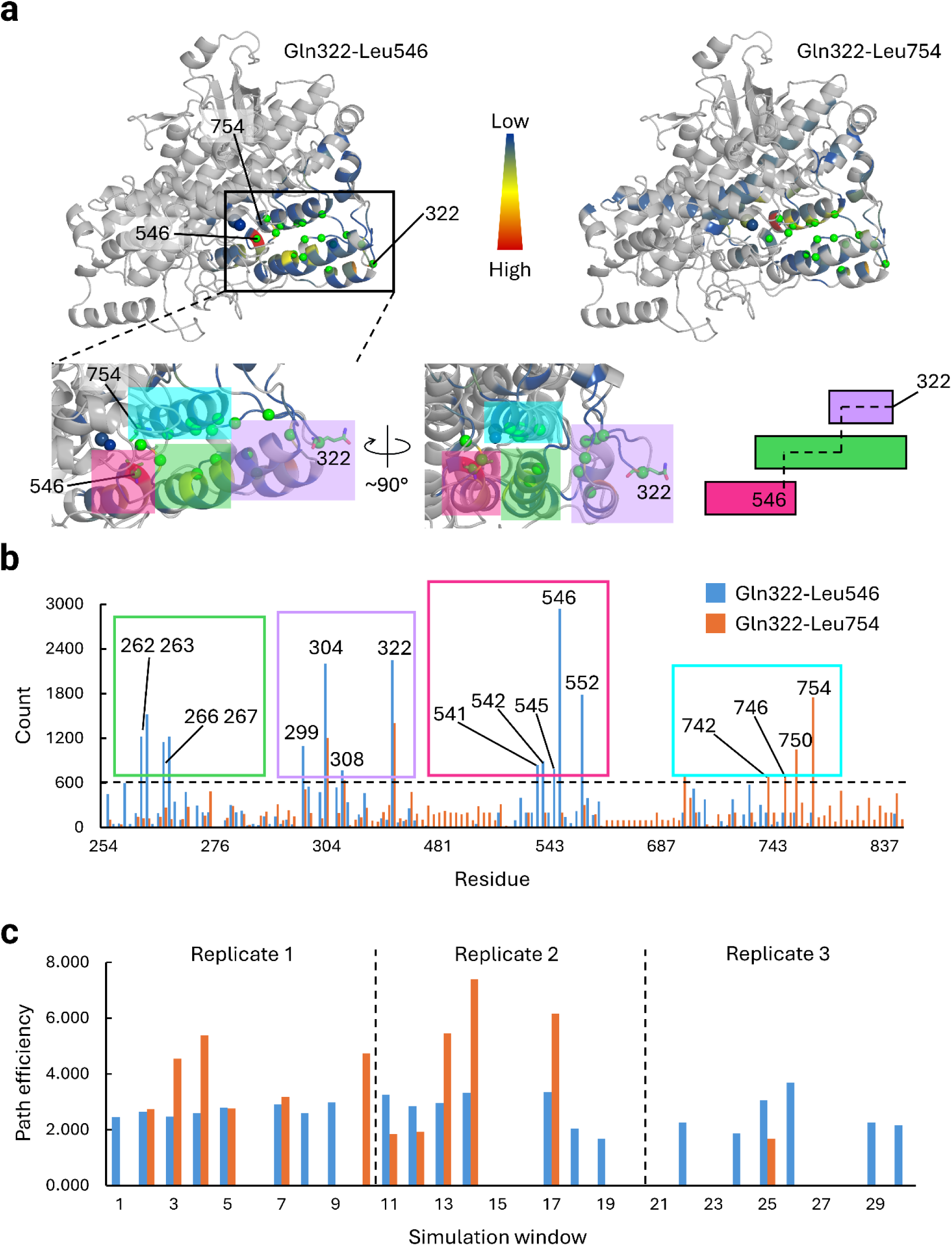
Linear mutual information (LMI)-based network comparison for the catalytic domain (residues 147-839) of SLO, refined by distance constraints for the Gln322–Leu546 and Gln322–Leu754 pathways. (a) Residue occurrence, color-coded from blue (minimum frequency), yellow (medium frequency), to red (maximum frequency). The active site Fe(II) and bound oxygen are shown as spheres that are colored based on occurrence, in the same fashion as amino acid residues, because they are part of the LMI-based network. Representative experimentally reported network residues (Leu262, Phe270, Leu299, Ile307, Ile313, Leu316, Tyr317, Gln322, Leu325, Leu546, Ile552, Ile553, Leu742, Ile746, Ser749, and Val750)^7,23^ are represented by green C_α_ spheres. The C_α_ positions of Leu546, Leu754, and Gln322 are pointed and labeled. The inset shows the enlarged region between Gln322 and the active site. Active site residues Leu546 and Leu754 and surface residue Gln322 are shown in green sticks. The color shades are used to distinguish four helical regions. An inset of the same region in an approximately 90° rotated view is provided to show the staggered side-by-side placement of the multiple helices. A simplified cartoon on the right shows the same placement of these three helices with the same color. The dashed line represents the approximate residue pathway connecting Gln322 and Leu546. (b) Column chart of residue occurrences for Gln322–Leu546 (blue) and Gln322–Leu754 (orange). Only residues with nonzero occurrences are included. Using the dashed line as a cutoff (>600 counts over the full 2 µs), key residues are identified and grouped into four sets of peaks with the same color scheme as the inset in Figure 3a. (c) Column chart of top path efficiencies for Gln322–Leu546 (blue) and Gln322–Leu754 (orange) across simulation windows. The path efficiency is calculated as the sum of the logarithm-converted LMI values along the path: ∑ − log(*LMI*). As such, a lower value means a more efficient path. Windows 1–10, 11–20, and 21–30 correspond to replicates 1, 2, and 3, respectively.

Beyond an enhanced overall frequency, the Gln322–Leu546 pathway is localized to three contiguous helical regions (Figure 3a, left, inset). Residues Leu299, Ile304, and Ile308, located near Gln322, form the entry point for energy transfer. Residues Leu262, Ser263, Val266, and Gln267 define an intermediate segment, while Leu541, Ala542, Ser545, and Ile552 form the terminal region proximal to Leu546. These residues are in similar positions but in some instances numerically different from those in the final experimental conduit.^7,23^ This indicates that computational modeling augments experimental results by revealing complementary side chains and pathways accessible within the simulated ensemble. The identified regions overlap with those previously defined through experiment and are shown here to correspond to three parallel helices, through which communication proceeds in order: Gln322 → helix 1 → helix 2 → helix 3 → Leu546 (Figure 3a, cartoon). Within each helix, correlated motion followed the i/i±4 residue pairs characteristic of α-helical hydrogen bonding (e.g., Ile304/Ile308, Leu262/Val266, Ser263/Gln267, and Leu541/Ser545). Inter-helix connectivity will be mediated through side-chain contacts.

The Gln322–Leu754 pathway is more diffuse, with residue involvement broadly distributed and lacking distinct clusters (Figure 3a, right). The only prominent segment is a helix adjacent to Leu754, where residues Leu742, Ile746, and Val750 show elevated occurrences, consistent with the same i/i±4 α-helical pattern described above (Figure 3a, left, insert).

We next compared the most probable communication paths across simulation windows, as these represent the highest-likelihood routes for energy transfer. Path efficiency, *Eff*, was quantified by the cumulative –log(LMI) of all edges along a path, such that lower values indicate stronger correlations and fewer residues involved. For Gln322–Leu546, a valid path was identified in 21 of 30 simulation windows, compared to only 12 of 30 for Gln322–Leu754 (Table S2). The mean path efficiency for Gln322–Leu546 was 2.679, compared to 3.988 for Gln322–Leu754, indicating stronger dynamic connectivity as shown in Figure 3c.

Direct comparison of the two paths (Leu 546 and Leu 754) within each window further emphasized this trend. In 10 windows, only a Gln322–Leu546 path was present, whereas only one window yielded exclusively Gln322–Leu754, with a path efficiency of 4.746—among the weakest communication events observed. In the 11 windows where both paths could be identified, Gln322–Leu546 was significantly better in 5 cases (Δ*Eff* = *Eff*_Gln322−Leu546_ − *Eff*_Gln322−Leu754_ ≤ −2), marginally different in 3 (|Δ*Eff*| < 0.5), and longer in 3 cases (2 > Δ*Eff* ≥ 0.5), though none showed a significant advantage for Gln322–Leu754 (Δ*Eff* ≥ 2).

In the remaining 8 windows, no paths were found for either active site-surface residue pair, largely within a single replicate. It is expected that thermal equilibration throughout SLO will be complete before the 200 ns cutoff (Figure S1b). The observed variability of each analyzed window is, thus, a consequence of the highly stochastic nature of SLO thermal energy activation. This property highlights the importance of extensive sampling to capture structure-based, protein scaffold-encoded energy transfer networks.

### Exploration of alternative protein surface loops as sites for thermal energy transport from solvent to active site residues in SLO

To examine whether the observed distal preference for communication between the protein surface Gln322 and active site Leu546 might be extended to other surface loop regions of SLO, we evaluated a range of potential alternative residues on the protein-solvent interface. Residues were manually selected from the 3PZW crystal structure^9^ using three criteria: (i) side chains exposed to solvent, (ii) location on a surface loop (assigned in *PyMOL*^24^), and (iii) exclusion of nonpolar residues whose side chains only contain C and H atoms (Gly, Ala, Val, Leu, Ile, Pro, Phe). The latter residues can be in the loop, but were not selected as the residues to start the network search. This yielded 73 residues across 12 loops (Figure 4a and Table S3).

**Figure 4.**
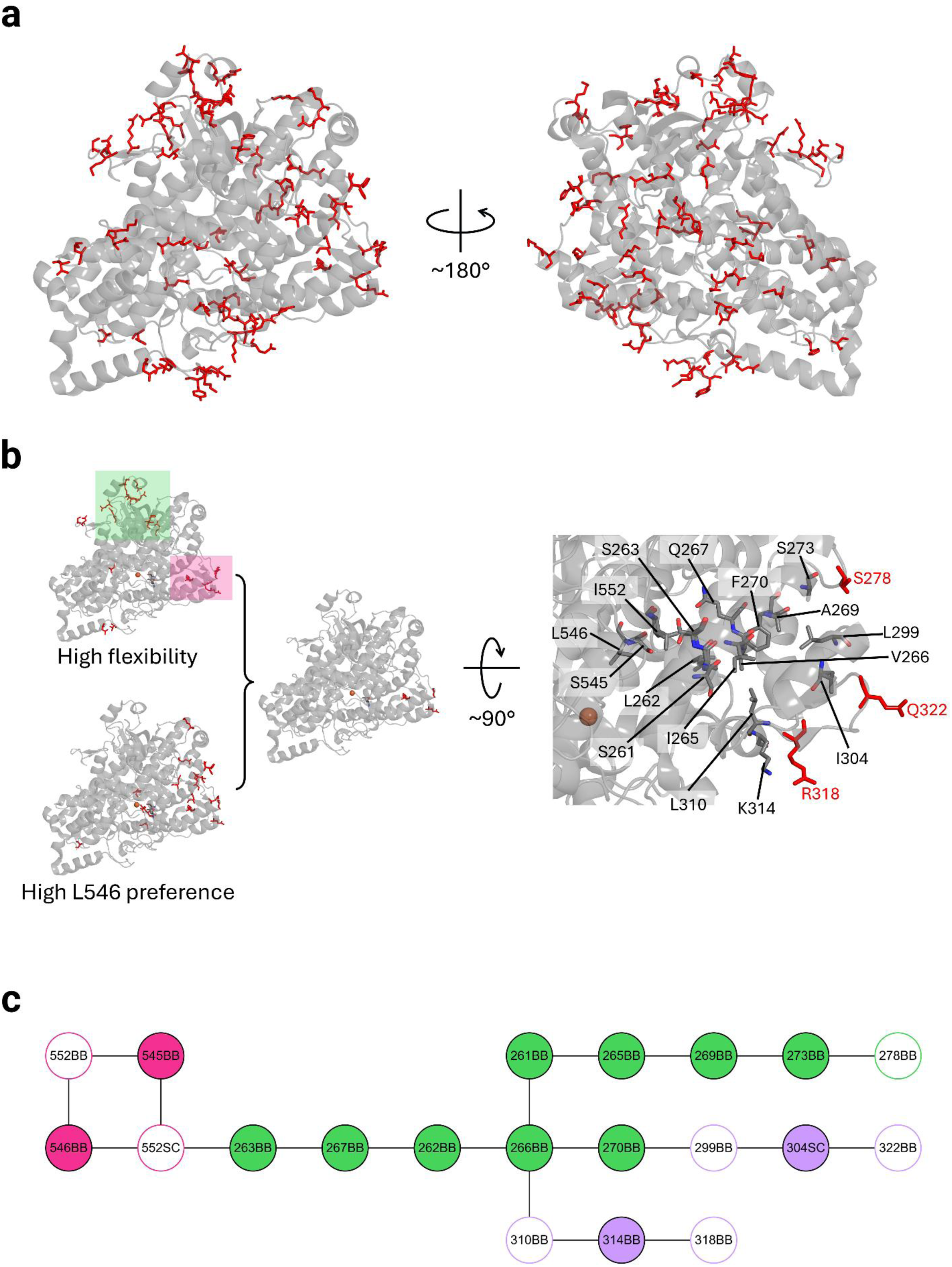
Surface-initiated dynamic communication converges on a specific region within the enzyme to channel energy to the active site. (a) Spatial distribution of all selected solvent-exposed loop residues shown as red sticks. (b) Cross-comparison of flexibility and Leu546 preference recovers the experimentally reported region. Left: top 20 residues with the highest flexibility (assessed by residue RMSF of backbone C, C_α_, and N atoms) or Leu546 preference (identified by LMI-based network analysis) shown as red sticks (See Table S3). Flexible regions are highlighted in pink and green. Right: zoomed-in view of the region formed by residues connecting the three recovered surface residues (Ser278, Gln322, and Arg318) to Leu546, with the corresponding residues in the best paths shown in sticks. We exclude the loop residues whose path occurrences towards both Leu546 and Leu754 are less than 1000, such as Asp320. The structures in a and b are based on crystal structure 3PZW, with only the catalytic domain (His147–Ile839) represented in cartoon. (c) The residue-node paths connecting Ser278, Gln322, and Arg318 to Leu546 are illustrated, where BB and SC represent backbone and side-chain nodes, respectively. For a given source, neighbor searches were initiated from either the backbone or side-chain node. Only nodes involved in the top 1 (out of all simulation replicates and windows) path connecting each pair of terminal residues are shown. The nodes are color-coded according to the corresponding regions highlighted in Figure 3a (inset) and Figure 3b, with identical colors used for matching regions. The computationally identified amino acids overlap significantly with the experimentally implicated regions of SLO (residues 263-299, 304-322, and 546-553). Nodes with solid fills represent residues within helices, whereas nodes outlined in color represent residues in loops adjacent to those helices.

Flexibility was assessed by residue-wise RMSF of backbone C, C_α_, and N atoms. The most flexible residues fell into two regions: residues Arg277, Ser278, Arg318, Asp320, and Gln322 on the previously identified initiation loops^7^ (highlighted in pink in Figure 4b), and residues Glu363, Asp391, Tyr393, Arg438, Glu439, Asp440, Asp594, Ser596, and Glu606 on a second surface region prominent in the catalytic domain (highlighted in green in Figure 4b). Notably, this second region contains Ser596, an experimental negative control for energy transfer.^25^ Both regions are enriched in ionic residues (Glu, Asp, Arg), consistent with their strong solvent interactions.

RMSF alone (Figure S3 and Table S3), however, overestimates potential initiation sites, including residues such as Ser596 showing no effective transfer. Using our LMI-based network protocol, we identified solvent-to-active site communication paths and counted their path occurrence across all node types, simulation windows, and replicas (maximum 6000 per residue). Among the 20 most flexible residues, 7—including Asp320, Thr174, and Ser596—exhibited fewer than 1/6 of maximum paths (i.e., fewer than 1000 paths) to the active site, with Ser596 showing none (Table S3). Thus, correlated-motion analysis is required to recover the experimentally validated initiation loop.

To quantify the destination preference for Leu546 over Leu754, we calculated the logarithm of the path occurrence ratio as the preference score, log(Occr_path_546_/Occr_path_754_), for each of the 73 solvent-exposed loop residues. Residues most biased toward Leu546 showed preference scores from 0.159 to 1.204 (actual Occr_path ratios > 1), and those favoring Leu754 ranged from –0.620 to –0.049 (ratios between 0 and 1). Besides Gln329, this distribution reveals a generally equal preference towards Leu546 and Leu754 as the communication endpoint (Figure S4).

Cross-comparison of flexibility and communication preference identified three residues—Gln322, Arg318, and Ser278—that fall within both the top 20 RMSF values and the top 20 Leu546-preference scores. Note that we excluded residues exhibiting fewer than 1/6 of maximum path occurrences (< 1000) to the active site, e.g., Asp320 and Asn172, even if they show a preference towards Leu546 or Leu754. All three residues of the cross preference are located on the experimentally validated initiation loop region (Figure 4b, right).^7^ The residue composition of these paths (Figure 4c) shows that Gln322, Arg318, and Ser278 converge at Val266 before proceeding along a common route to Leu546. The involved regions overlap closely with those inferred from crystal structure analysis,^23^ placing these surface residues within the experimentally defined “cone” that dynamically communicates with the active site.

Four residues, His454, Glu439, Glu363, and Arg438, were recovered among the top 20 flexible residues, as well as the top 20 preference scores for Leu754. These reside in distinct surface regions dominated by antiparallel β-sheets (Figure S5a). All four residues show lower flexibility compared to Gln322, Arg318, and Ser278, although they are among the top 20 flexible residues. Note that we excluded residues showing fewer than 1/6 (1000) of the maximum path towards both L546 and L754 to exclude inert ones in thermal energy transfer. Path analysis further emphasized these differences. The best solvent–active site paths from Gln322, Arg318, and Ser278 to Leu546 had total –log(LMI) values of 1.676, 1.567, and 1.684, respectively, indicating strongly correlated dynamics. In comparison, His454, Glu439, Glu363, and Arg438 connected to Leu754 only through paths with considerably larger values: 2.674, 2.664, 2.902, and 2.556, reflecting weaker dynamic communication. Interestingly, by comparing the computed residue paths in SLO connecting protein-solvent interface and Leu754 to the experimentally defined thermal activation network in mammalian lipoxygenase 15-LOX-2,^26^ we found overlap between an SLO 754 residue path and a key peptide within the 15-LOX-2 thermal network (Figure S5b). This suggests that off-network routes in SLO present structurally conserved communication elements that could be utilized in evolved mammalian lipoxygenases that lack the unique thermal activation loop in SLO.

Together, these results provide a molecular framework for the experimental observation that Leu546 predominantly participates in the communication network: (i) energy transfer is initiated at highly flexible solvent-exposed loops, (ii) propagation follows pathways with efficient correlated dynamics, and (iii) the relative efficiency of these pathways determines which active-site residue is preferentially engaged. Looking beyond SLO, this analysis showcases a prediction protocol to identify protein surface sites for initiation of energy transfer from solvent to the reactive positions of enzyme-bound substrate(s); the latter is readily available from decades of detailed structural and mechanistic investigations on enzymes.

### Impact on Computed Networks of Mutations that Alter Activation Energies for k_cat_

To probe the effect of mutations on dynamic communication, we performed MD simulations on four variants with experimentally characterized differences in rates and activation energies: I553G, I553A, I552A, and L546A. These residues are identified either as in contact with the reactive bond of the substrate (Leu546, Figure 1), in remote contact with bound substrate (Ile553),^27^ or in contact with L546 itself (Ile552, Figure 4b), thus directly testing how structural perturbations reshape the communication pathways.

Using the same LMI-based analysis, we quantified total residue occurrences in correlated-motion networks connecting Gln322 with Leu546 or Leu754. In the wild type, Gln322–Leu546 communication involves 33,596 occurrences, compared to 26,686 for Gln322–Leu754 (Figure 5). Mutations substantially reduced overall occurrences. For Gln322–Leu546, values ranged from 29,548 in I553G to only 11,777 in L546A. For Gln322–Leu754, the range was 18,357 (I553G) to 8,204 (I552A). These reductions likely represent a loosening in residue packing due to reductions in side chain size, which can be expected to diminish correlated motions.

**Figure 5.**
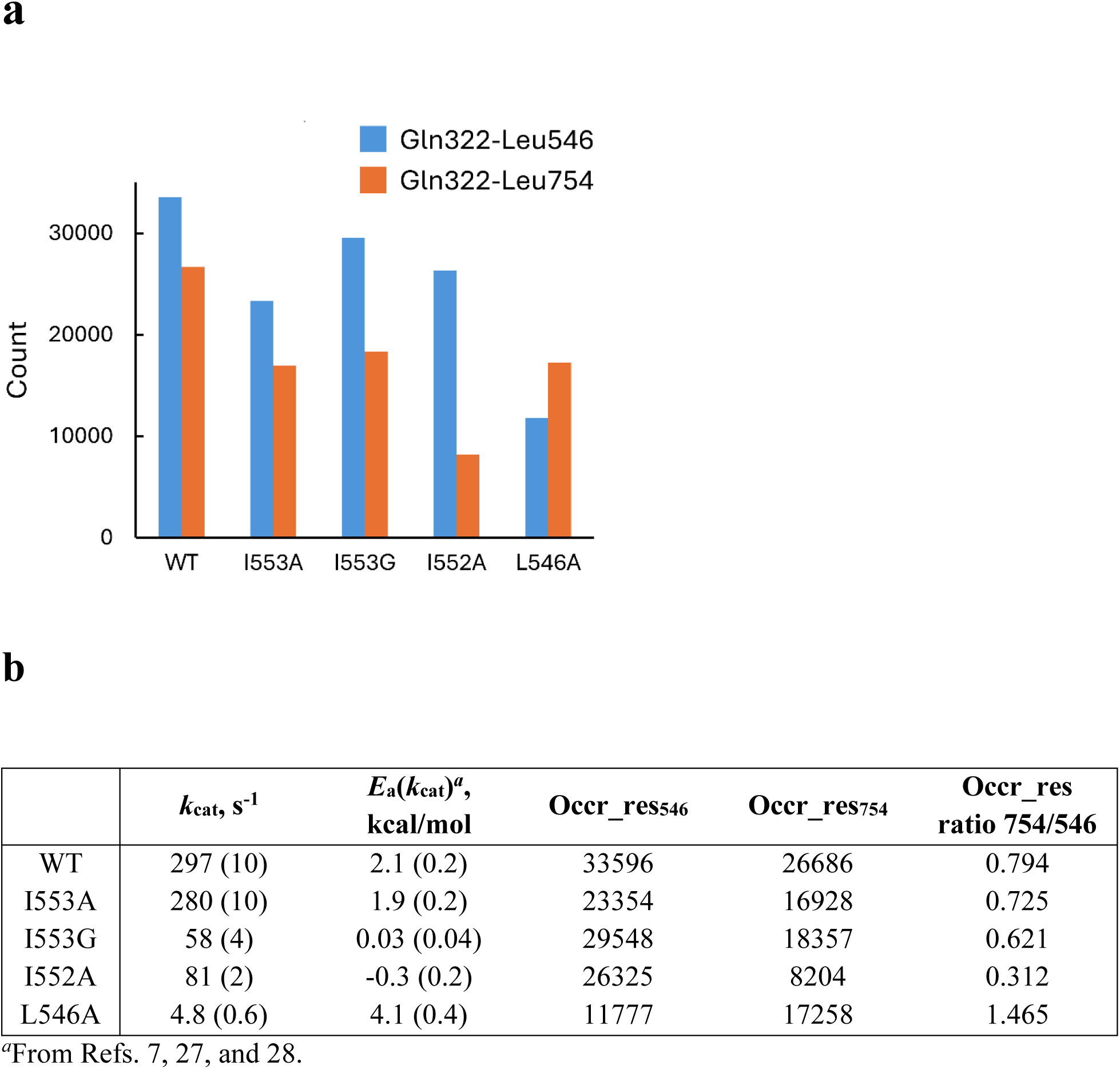
Mutation effects on the residue network. (a) Column chart of total residue occurrences in the Gln322–Leu546 (blue) and Gln322–Leu754 (orange) networks. (b) The total residue occurrences for all soybean lipoxygenase variants are compared to the *k_cat_* and *E*_a_(*k*_cat_) data for C-H activation from previous literature.^7,27,28^ The residue occurrence ratio is calculated as 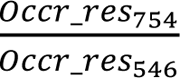.

The extent of disruption differs sharply between the two networks (Figure 5a). For instance, in I552A, the Gln322–Leu546 network decreased by 1.28-fold relative to WT, whereas the Gln322–Leu754 network decreased 3.25-fold. By contrast, reducing the side chain size at L546 with the L546A mutation reverses the preference between the two networks to favor L754. This supports the view of multiple potential pathways inherent within a protein structure and is consistent with a completely different thermal energy network for the human lipoxygenase 15-LOX-2 (Figure S5) that lacks the distinct loop structure of SLO and is characterized by an elevated activation energy of 6.8 kcal/mol (arachidonic acid, native substrate)^26^ compared to 2.1 kcal/mol for SLO above (linoleic acid, native substrate).^28^

To quantify further the network preference as a function of mutation, we consider the residue occurrence ratio Occr_res_754_/Occr_res_546_ in comparison to experimentally observed *E*_a_(*k*_cat_) and *k*_cat_ values shown in Figure 5b. Position 553 interacts with a remote position of the substrate and has been shown to influence the pathway for the binding of linoleic acid to SLO.^29^ In this context, I553A serves as a control, given its near identity of *E*_a_(*k*_cat_) and *k*_cat_ to WT, as well as an almost unchanged value for Occr_res_754_/Occr_res_546_. In the case of I553G and I552A, which have been shown from hydrogen deuterium exchange experiments and RT X-ray studies to perturb the thermal network of SLO,^5,7^ there are noticeable changes to the occurrence ratio. While the I553 was not identified using LMI-based network analysis, there is a detectable decrease in Occr_res_754_/Occr_res_546_. This becomes more significant for I552A, with a surprising direction that suggests preferential stabilization of the Gln322-Leu546 pathway. We note from previous studies that I553G and I552A lead to decreased experimental values of *E*_a_(*k*_cat_) that approach zero.^7,27^ This behavior can be rationalized by the introduction of mutant-induced, off-path equilibrated structural changes that are entropically favored/enthalpically disfavored relative to the native enzyme structure. Such alternate ground state structures may be expected to impede the flow of thermal energy from the protein-solvent interface to the active site, requiring an enthalpy-driven reversal to native-like configurations before productive heat flow. The net effect of such behavior would be to mask the contribution of the intrinsic uphill enthalpic barrier for C-H bond cleavage. One of the most important observations from Figure 5 is that L546A reverses the pathway choice, in full support of a keystone role for Leu 546 in the construction of the primary thermal network (Gln322-Leu546).

## DISCUSSION

The presented LMI-based network protocol (Figure 2) can recapitulate the major features of the experimentally observed distal communication in soybean lipoxygenase. Our protocol targets two experimentally identified active site residues that reside on opposite sides of the reactive carbon of bound substrate^8,25^ as putative active site nodes for thermal energy transfer. The specificity of the corresponding solvent-exposed node has been surveyed using RMSF analyses of multiple loops around the protein surface (Figure 4). The aggregate simulations recover the experimentally identified cone of residues connecting Gln322 at the surface to Leu546 in the active site, finding this network to be favored in relation to an alternate Gln322–Leu754 network.^7,23^ The agreement with experiment extends to the negative control residue Ser596, which displays high flexibility but fails to initiate productive communication in both experiment^25^ and computation. These results highlight the necessity of correlation-based network analysis, rather than residue-wise flexibility alone, for identifying functional communication pathways.

The simulations further reveal why Leu546 is favored over Leu754: pathways to Leu546 are shorter, more correlated, and more localized, converging through a set of three helices that act as an efficient conduit (Figure 3a, left). In contrast, Leu754 is reached only through longer, weaker, and more diffuse paths. The computationally derived mechanism explains the experimental distinction between in-network and off-network residues,^30^ and clarifies the structural basis for anisotropic communication in SLO.

The preference for Leu546-directed communication is in full agreement with experimental findings that mutation of Leu546 (L546A) alters the activation barrier for hydrogen-deuterium exchange that arises from changes in local protein unfolding at the protein surface, *E*_a_*(k*_HDX_), whereas mutation of Leu754 (L754A) does not.^7,23^ As previously discussed, the Gln322–Leu546 pathway appears ideally positioned to provide a thermally initiated optimization of the two primary components of the reorganization barrier to C-H activation: The H-donor and acceptor distances and the electrostatic optimization of reactant-product vibrational mode overlap.^4^

Computational analysis of experimentally assessed sites of mutation further strengthens the analysis. The directional change in the occurrence ratio, Occr_res_754_/Occr_res_546,_ upon mutation of Leu546 corroborates its key role in directing remote thermal energy from solvent to the reactive position of substrate, reinforcing emerging molecular models for the construction of long-range thermal energy transport pathways in SLO.^4^ The obtained results do not rule out an interactive role between Leu546 and Leu754 in mediating the reaction activation energy, and this aspect will require further investigation into other related mutations.

More broadly, computational analysis of SLO expands our emerging understanding of reaction-specific, distal thermal networks.^4,31,32^ Unlike allosteric regulation,^33–35^ which redistributes pre-existing conformational ensembles, these networks represent anisotropic pathways of energy flow that couple solvent collisions to chemical steps. By unifying residue-level dynamics with experimental observables, our approach provides a possible generalizable computational strategy for mapping intra-enzyme communication and connecting it to catalysis.

## CONCLUSIONS

The application of long-timescale MD simulations with LMI-based network analysis, which has been refined by distance constraints, can reproduce and extend experimental findings in SLO. This work introduces a computational tool for the corroboration of experimentally reported thermal activation networks as well as for the prediction of the large repertoire of as yet uncharacterized thermal energy networks among enzyme catalysts. The combination of experimental and computational advances supports a shift from enzyme models that rely on static protein structures to the involvement of dynamical scaffolds that enable site-specific thermal energy flow from solvent to active site.

## ACKNOWLEDGMENTS

This work was supported by grants from the NIH (GM118117) and NSF (2231081) to J.P.K. This research used the Savio computational cluster resource provided by the Berkeley Research Computing program at the University of California, Berkeley (supported by the UC Berkeley Chancellor, Vice Chancellor for Research, and Chief Information Officer). Y.J. thanks Dr. Nigel Richards, Dr. Susan Miller, and Professor Zhongyue Yang for their assistance in enzyme structure preparation and inspiring discussions.

## SUPPORTING INFORMATION

Detailed computational methods; root mean square deviation of atomic positions and temperature for the MD simulation of soybean lipoxygenase; example linear mutual information matrix calculated from molecular dynamics simulations; root mean square fluctuation of the catalytic domain of soybean lipoxygenase; active site residue preference for the selected surface loop residues; cross-comparison of flexibility and Leu754 preference of the selected solvent-exposed residues leads to a different region compared to the experiment; occurrence of residues within the correlated-motion-based network of the soybean lipoxygenase; efficiency of the best path connecting Gln322 and Leu546 or Gln322 and Leu754 in each simulation window; solvent-exposed residues’ proxies related to recovering the experimental initiation loop and Leu546-directed cone. (PDF)

## AUTHOR INFORMATION

### NOTES

The authors declare no competing financial interest.

