## Supporting Information for "Correlated Motion-Based Residue Network Analysis Reveals the Distal Thermal Activation in Soybean Lipoxygenase"

**Contents**

**Text S1.** Detailed computational methods. Page S2
**Figure S1.** Root mean square deviation of atomic positions and temperature for the MD simulation of soybean lipoxygenase. Page S3
**Figure S2.** Example linear mutual information matrix calculated from molecular dynamics simulations. Page S4
**Figure S3.** Root mean square fluctuation of the catalytic domain of soybean lipoxygenase. Page S4
**Figure S4.** Active site residue preference for the selected surface loop residues. Page S5
**Figure S5.** Cross-comparison of flexibility and Leu754 preference of the selected solvent-exposed residues leads to a different region compared to the experiment. Page S6
**Table S1.** Occurrence of residues within the correlated-motion-based network of the soybean lipoxygenase. Page S7
**Table S2.** Efficiency of the best path connecting Gln322 and Leu546 or Gln322 and Leu754 in each simulation window. Page S9
**Table S3.** Solvent-exposed residues' proxies related to recovering the experimental initiation loop and Leu546-directed cone. Page S10

**Text S1.** Detailed computational methods.

*Molecular Dynamics Simulations.* The initial structure was based on the crystal structure of soybean lipoxygenase (PDB ID 3PZW).[1] Unmodeled loops (Met1–His6, Glu21–Asp30, Ile117–Gln120) were added from another crystal structure 1YGE.[2] The Fe(II)-bound groups (His499, His504, His690, Ile839, and one water molecule) were parameterized with the Metal Center Parameter Builder (*MCPB.py*).[3] Using Fe(II) bound to $H_2O$ or Fe(III) bound to $OH^-$ is unlikely to affect the simulation outcome significantly. No substrate or intermediate was included in the active site. The docked pose of the substrate linoleate remains uncertain, as its carboxylate group may adopt either an "in-facing" orientation (Figure 1) or an "out-facing" configuration corresponding to a head-tail reversal. Mutations were introduced by changing the residue name and removing side-chain atoms except $C_\beta$ in the wild-type coordinate file; missing residues were then added in *tleap*. The solvated truncated octahedron box ensured a minimum distance of at least 10 Å between the protein and box boundaries, containing ~32,000 water molecules, with sodium ions added to neutralize the system.

Energy minimization proceeded stepwise from the Fe(II)-bound water to the entire system. The box was heated from 0 to 298.15 K over 36 ps under constant volume, followed by a 4 ps equilibration at 298.15 K (constant volume), and then a 1 ns equilibration at 298.15 K and constant pressure, 1 atm. Backbone atoms (C, $C_\alpha$, N) were restrained with 2.0 kcal·mol$^{-1}$·Å$^{-2}$ during these steps. A second 10 ns equilibration at 298.15 K and 1 atm was run without restraints, followed by a 4 μs production run. Snapshots were saved every 50 ps, yielding 80,000 frames. Three independent simulations were performed per enzyme variant.

*Network Identification Protocol.* To minimize interference from interdomain motion, the N-domain (Met1–Asn146) was excluded, and only the catalytic domain (His147–Ile839) was analyzed. RMSD analysis showed that the catalytic domain equilibrated after ~2 μs (Figure S1), so analyses focused on the last 2 μs of the trajectory (2–4 μs). This segment was divided into 10 equal windows, each containing 4000 snapshots.

The original trajectory was converted into a node trajectory using the *vector* command of *cpptraj*.[4] Each residue was represented by two nodes: backbone (BB; $C_\alpha$, C, O, N atoms) and side chain (SC; heavy atoms), except glycine, which lacked an SC node. Node coordinates were defined by the centers of mass of the corresponding atoms. Fe(II) and the oxygen of its bound water were included as two additional nodes. In total, the network contained 1349 nodes, except for variant I553G, which contained 1348.

Pairwise linear mutual information (LMI) was calculated between all nodes, resulting in a symmetric 1349 × 1349 matrix using the Python package *correlationplus*.[5] To eliminate local correlations dominated by covalent bonds, entries for residues separated by fewer than four sequence positions were removed.

Neighbor searches were initiated from either the backbone or side-chain node of a given source residue. A neighboring node was defined as having an LMI ≥ 0.40 with the source node. An *n*th-level neighbor required *n* steps to connect to the source node. Results from BB- and SC-initiated searches were combined to determine the neighbor residues of the source. A residue could therefore be classified as more than one type of neighbor. The search proceeded until the target residue was reached or until the 10th level.

If the target was identified at the 0.40 cutoff, the search was repeated at higher thresholds (0.45, 0.50, etc.), incrementing by 0.05, until the target could no longer be found. The highest cutoff at which the target was still reached was recorded as the effective LMI cutoff between the source and target. The source nodes and all nodes along the path to the target defined the **initial subnetwork**.

Contact distances between all residue pairs in the initial subnetwork were calculated with the *nativecontacts* command in *cpptraj*,[4] defined as the minimum heavy-atom distance between residues. The following criteria then filtered the LMI matrix:

1. Residues belonged to the initial subnetwork (all combinations of source/terminal BB/BB, BB/SC, SC/BB, SC/SC considered).
2. LMI ≥ cutoff.
3. ≥ 75% of sampled snapshots had contact distances ≤ 4.5 Å.[6]

The resulting network was defined as the **refined subnetwork**. For weighting, converted LMI (*cLMI*) values were calculated as:

$$cLMI = -\log(LMI)$$

This conversion serves two purposes. First, because LMI represents a "larger-is-better" metric of correlation, whereas path-finding algorithms interpret lower edge weights as more favorable, the sign must be inverted to make stronger correlations correspond to shorter paths. Second, applying the logarithm further accentuates this weighting: when LMI = 1 (perfect correlation), the converted value *cLMI* = 0, indicating no penalty for movement between the corresponding nodes. Conversely, as LMI approaches 0 (minimal correlation), *cLMI* increases toward infinity, imposing a very high penalty on weakly correlated connections.

Yen's algorithm[7] was then applied to identify the top 50 communication paths, using the shortest path determined by Dijkstra's algorithm[8] as the reference, by minimizing the total *cLMI* between the source and target residues. Analyses were performed separately for source/target node combinations BB/BB, BB/SC, SC/BB, and SC/SC. This separation was only applied to source and target nodes. The resulting paths, including their nodes and edges, defined the **final subnetwork**.

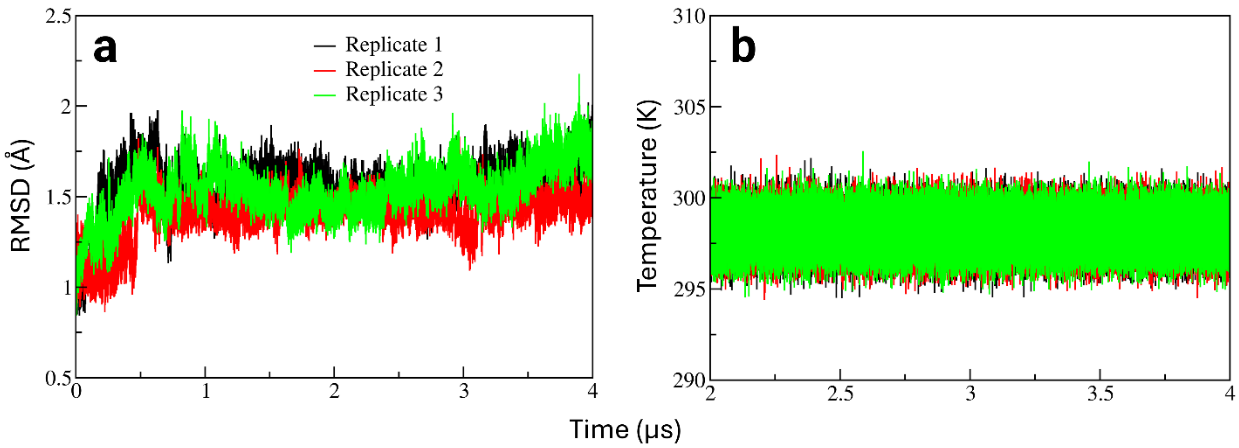


**Figure S1.** Root mean square deviation (RMSD) of atomic positions (a) and temperature (b) for the molecular dynamics simulation of soybean lipoxygenase. (a) Backbone C, $C_\alpha$, and N atoms are included to calculate RMSD. The reference is the initial structure before energy minimization to enable comparisons between replicates. Only the catalytic domain (His147–Ile839) is considered to eliminate the interference of inter-domain movements. (b) All atoms in the simulation box are included to calculate the temperature. The average temperatures for replicates 1 to 3 are 298.16 ± 0.89, 298.17 ± 0.89, and 298.17 ± 0.89 K, respectively. Only the result of the second 2 μs simulation (used for network analysis) is shown.

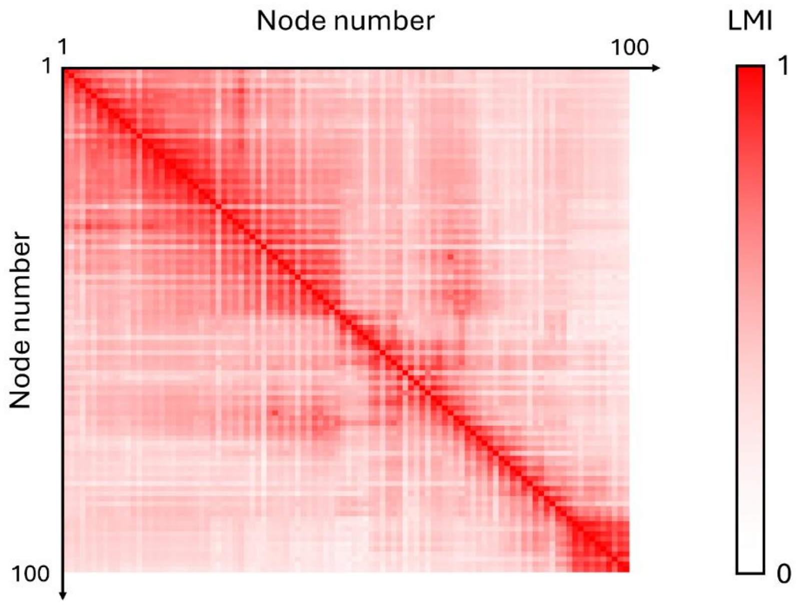


**Figure S2.** Example linear mutual information (LMI) matrix calculated from molecular dynamics simulations. This matrix contains the first 100 nodes, Asn146 backbone to Ser198 side chain. The full-size matrix is 1349 × 1349, containing nodes from Asn146 backbone to Ile839 side chain, Fe(II), and its bound water oxygen. The LMI values are color-coded between white (0) and red (1).

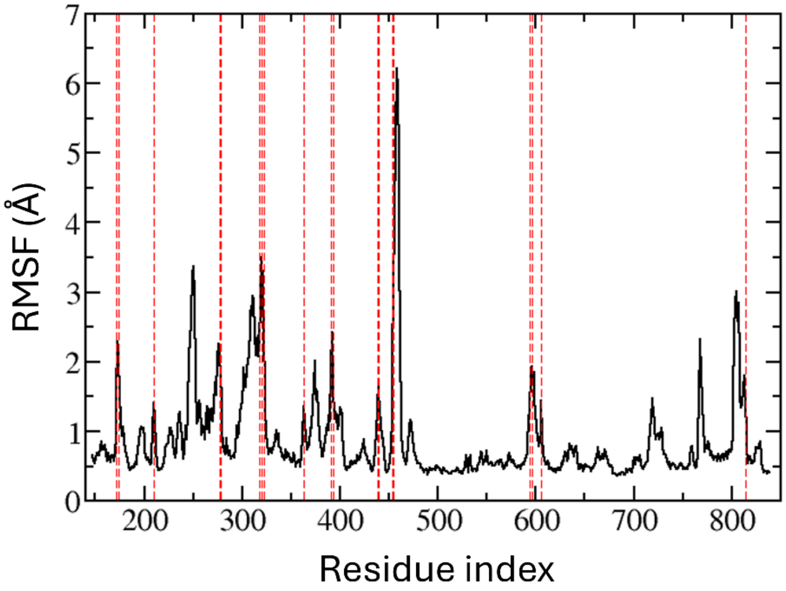


**Figure S3.** Root mean square fluctuation (RMSF) of the catalytic domain of soybean lipoxygenase. The RMSF is averaged by residue. Only backbone C, $C_\alpha$, and N are included. The red dashed lines indicate the top 20 residues with the highest RMSF value among the 73 selected solvent-exposed ones shown in Table S3: Asn172, Thr174, Ser210, Lys277, Ser278, Arg318, Asp320, Gln322, Glu363, Asp391, Tyr393, Arg438, Glu439, Asp440, His454, Ser455, Asp594, Ser596, Glu606, Gln814.

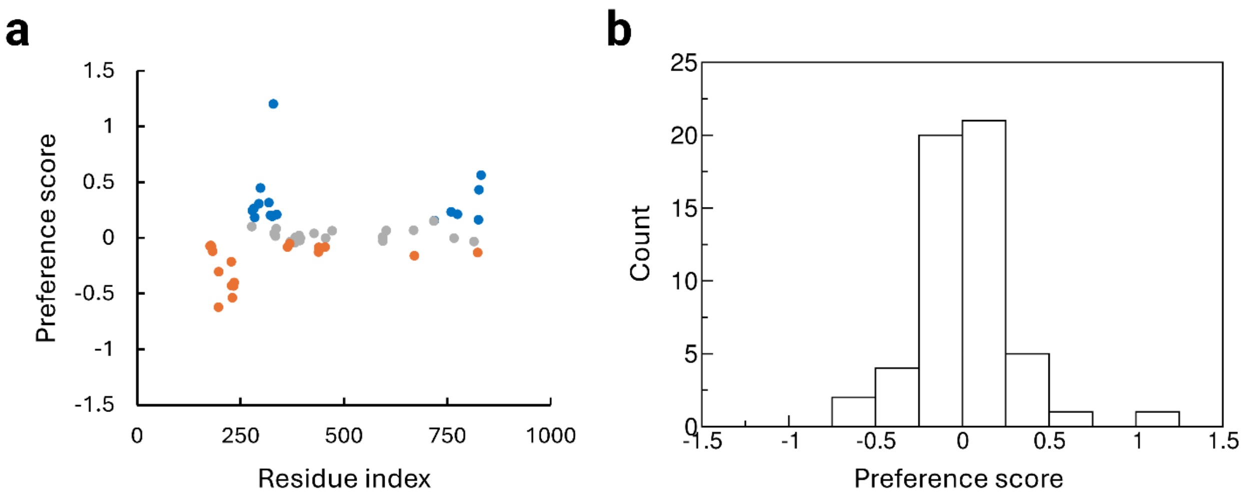


**Figure S4.** Active site residue preference for the selected surface loop residues. The preference score is calculated as log($Occr\_path_{546}$/$Occr\_path_{754}$), where Occr_path represents the path occurrence of a specific loop residue towards active site residues Leu546 or Leu754. (a) Scatter plot of the preference score for each loop residue. The horizontal axis represents the index of the loop residue. The top 20 loop residues with a high Leu546 preference (blue), top 20 with high Leu754 preference (orange), and others with trivial preference (gray). (b) Histogram of the preference score shown in (a), where positive values are 546-preferred, and negative values are 754-preferred. The histogram bin size is 0.25. Both plots exclude the loop residues whose path occurrences toward Leu546 and Leu754 are both less than 1000. The preference score data corresponds to those shown in Table S3.

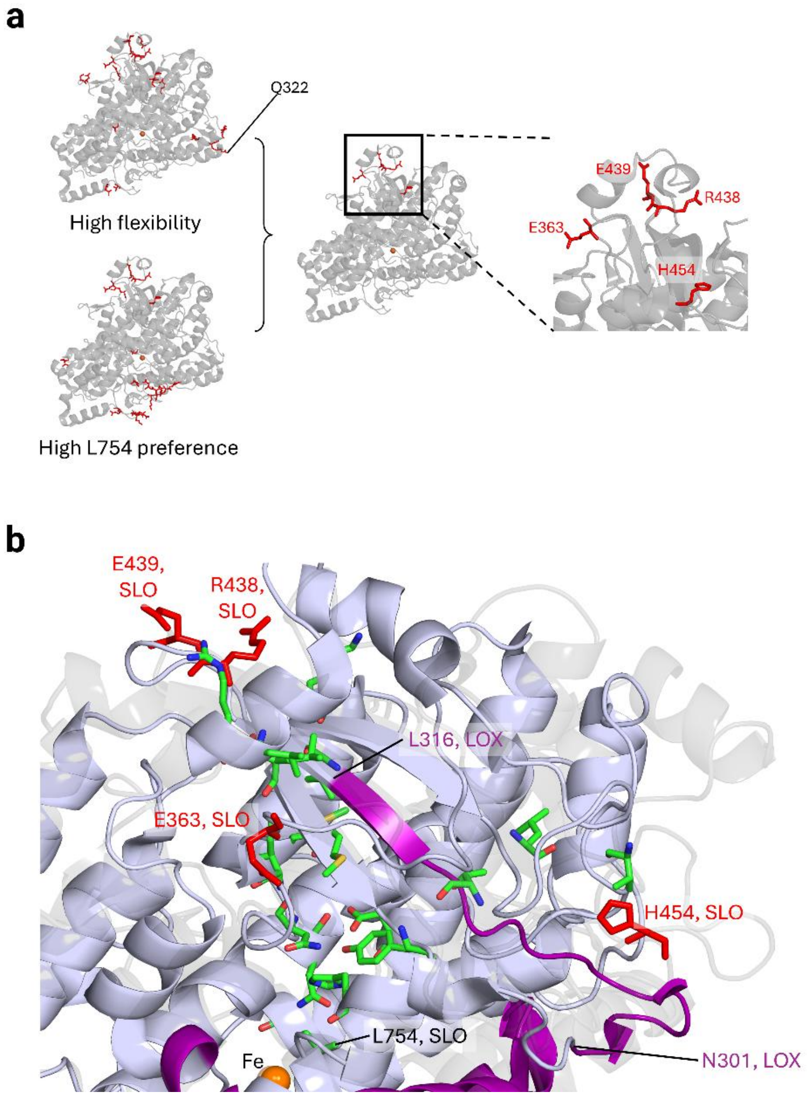


**Figure S5.** Cross-comparison of flexibility and Leu754 preference of the selected solvent-exposed residues leads to a different region compared to the experiment. (a) Left: top 20 residues with the highest flexibility (assessed by residue RMSF of backbone C, $C_\alpha$, and N atoms) or Leu754 preference (identified by LMI-based network analysis) shown as red sticks (See Table S3). As a reference, Gln322's position is labeled. Gln322 is among the top 20 most flexible residues, but not among the top 20 with the highest Leu754 preference. Right: zoomed-in view of the region from the cross-comparison, with the four residues with both high flexibility and Leu754 preference shown in red sticks. (b) Comparison of the best residue paths (green sticks) connecting Leu754-preferred surface residues (His454, Glu439, Glu363, and Arg438) to Leu754 in SLO and the thermal activation network (purple cartoon) in mammalian lipoxygenase 15-LOX-2.[9] The structure of SLO is shown as a transparent gray cartoon. The crystal structure of 15-LOX-2 (PDB ID: 4NRE[10]) is overlaid to SLO using the *align* command in *PyMOL*[11] and is shown as a light blue cartoon. The His454–Leu754 residue pathway in SLO crosses one of the key network peptides, Asn301–Leu316 (purple), in 15-LOX-2.[9] Other surface–Leu754 pathways cross the β-sheet adjacent to this network peptide.

**Table S1.** Occurrence of residues within the correlated-motion-based network of the WT soybean lipoxygenase connecting Gln322 and Leu546 or Gln322 and Leu754. The residues are listed in descending order of the corresponding occurrence. The bolded ones belong to those in the reported experimental network.[12,13] The sums of Gln322–Leu546 and Gln322–Leu754 residue occurrences are 33596 and 26686, respectively. As a reference, a maximum residue occurrence is 50 (top paths) × 4 (backbone and side chain node combinations) × 10 (simulation windows) × 3 (simulation replicates) = 6000. Two reasons account for the occurrence of terminal residues, Leu546, and Gln322, not being a multiple of 50: First, fewer than 50 paths can be found in some cases. Second, backbone (BB) and side chain (SC) nodes of the same residue may appear in one path. For example, in path 322BB → 304BB → … → 552BB → 546BB → 542BB → 546SC (intermediate residues between 304BB and 552BB are omitted), Leu546 backbone node (546BB) is an intermediate node to reach Leu546 side chain node (546SC) from Gln322 backbone node (322BB). This also accounts for the unequal occurrence of the source and target residues, even though they exist in each identified path.

| Index | Residue index | Gln322-Leu546 | Residue index | Gln322-Leu754 | Index | Residue index | Gln322-Leu546 | Residue index | Gln322-Leu754 |
|---|---|---|---|---|---|---|---|---|---|
| 1 | **546** | 2937 | 754 | 1750 | 61 | 319 | 102 | 273 | 157 |
| 2 | **322** | 2250 | **322** | 1402 | 62 | 278 | 100 | 286 | 157 |
| 3 | 304 | 2202 | 304 | 1203 | 63 | 534 | 100 | 297 | 150 |
| 4 | **552** | 1784 | **750** | 1050 | 64 | 326 | 96 | 261 | 146 |
| 5 | 263 | 1523 | 694 | 700 | 65 | 324 | 88 | 265 | 146 |
| 6 | **262** | 1225 | **742** | 679 | 66 | 731 | 85 | **313** | 138 |
| 7 | 267 | 1223 | **746** | 679 | 67 | 744 | 75 | 755 | 136 |
| 8 | 266 | 1148 | 308 | 635 | 68 | 740 | 73 | 278 | 128 |
| 9 | **299** | 1095 | **299** | 515 | 69 | 323 | 67 | 690 | 128 |
| 10 | 542 | 891 | 763 | 492 | 70 | 548 | 57 | **262** | 127 |
| 11 | 541 | 838 | 274 | 488 | 71 | 258 | 54 | 303 | 125 |
| 12 | 545 | 790 | **325** | 476 | 72 | 256 | 48 | 323 | 124 |
| 13 | 308 | 772 | 840 | 462 | 73 | 260 | 48 | 263 | 121 |
| 14 | 259 | 600 | 698 | 400 | 74 | 264 | 48 | 267 | 117 |
| 15 | 739 | 575 | 833 | 400 | 75 | 285 | 48 | 269 | 114 |
| 16 | 300 | 552 | 751 | 336 | 76 | 289 | 48 | 841 | 111 |
| 17 | **307** | 539 | 504 | 314 | 77 | 293 | 48 | 327 | 108 |
| 18 | 709 | 523 | 292 | 313 | 78 | 499 | 46 | 764 | 108 |
| 19 | **270** | 476 | 319 | 302 | 79 | 297 | 45 | 324 | 106 |
| 20 | 303 | 475 | **552** | 301 | 80 | 743 | 38 | 834 | 106 |
| 21 | 314 | 464 | 328 | 300 | 81 | 286 | 31 | 255 | 104 |
| 22 | 255 | 452 | 757 | 300 | 82 | 734 | 31 | 312 | 104 |
| 23 | 537 | 400 | 827 | 300 | 83 | 284 | 28 | 687 | 102 |
| 24 | **553** | 395 | 835 | 294 | 84 | 328 | 23 | 293 | 100 |
| 25 | 735 | 381 | 279 | 292 | 85 | 698 | 23 | 498 | 100 |
| 26 | 712 | 379 | 499 | 286 | 86 | 747 | 23 | 503 | 100 |
| 27 | 557 | 351 | **270** | 280 | 87 | 751 | 23 | 636 | 100 |

| | | | | | | | | | |
|---|---|---|---|---|---|---|---|---|---|
| 28 | 268 | 348 | 266 | 271 | 88 | 306 | 17 | 671 | 100 |
| 29 | 310 | 339 | 314 | 264 | 89 | **317** | 16 | 672 | 100 |
| 30 | 741 | 304 | 296 | 235 | 90 | 276 | 12 | 675 | 100 |
| 31 | 279 | 302 | 485 | 222 | 91 | 298 | 6 | 676 | 100 |
| 32 | 272 | 300 | 288 | 213 | 92 | 254 | 4 | 679 | 100 |
| 33 | **325** | 256 | 318 | 210 | 93 | 277 | 4 | 680 | 100 |
| 34 | 283 | 226 | 481 | 200 | 94 | 312 | 4 | 683 | 100 |
| 35 | 551 | 220 | 489 | 200 | 95 | 271 | 2 | 684 | 100 |
| 36 | 710 | 210 | 493 | 200 | 96 | 302 | 2 | 689 | 100 |
| 37 | 273 | 206 | 497 | 200 | 97 | 841 | 2 | 693 | 100 |
| 38 | 274 | 206 | 500 | 200 | 98 | | | 821 | 100 |
| 39 | 282 | 204 | 501 | 200 | 99 | | | 831 | 100 |
| 40 | **742** | 202 | 538 | 200 | 100 | | | **316** | 92 |
| 41 | 538 | 201 | 542 | 200 | 101 | | | 762 | 82 |
| 42 | 504 | 200 | **546** | 200 | 102 | | | **317** | 78 |
| 43 | 539 | 200 | 686 | 200 | 103 | | | 752 | 64 |
| 44 | 543 | 200 | 709 | 200 | 104 | | | 712 | 43 |
| 45 | 694 | 200 | 735 | 200 | 105 | | | 258 | 42 |
| 46 | **746** | 200 | 739 | 200 | 106 | | | 284 | 35 |
| 47 | **750** | 200 | 743 | 200 | 107 | | | 310 | 32 |
| 48 | 754 | 200 | 747 | 200 | 108 | | | 505 | 22 |
| 49 | 261 | 193 | 837 | 200 | 109 | | | 713 | 22 |
| 50 | 265 | 192 | 839 | 200 | 110 | | | 737 | 22 |
| 51 | 840 | 184 | **307** | 197 | 111 | | | 738 | 21 |
| 52 | **313** | 175 | 300 | 195 | 112 | | | **553** | 20 |
| 53 | 738 | 175 | 326 | 184 | 113 | | | 557 | 20 |
| 54 | 556 | 172 | 283 | 181 | 114 | | | 741 | 19 |
| 55 | 288 | 154 | 556 | 180 | 115 | | | 531 | 16 |
| 56 | 292 | 148 | 477 | 178 | 116 | | | 731 | 8 |
| 57 | 547 | 143 | 496 | 178 | 117 | | | 259 | 4 |
| 58 | 318 | 123 | 730 | 178 | 118 | | | 277 | 4 |
| 59 | 296 | 106 | 734 | 178 | | | | | |
| 60 | 269 | 102 | 756 | 164 | | | | | |

**Table S2.** Efficiency of the best path connecting Gln322 and Leu546 or Gln322 and Leu754 in each simulation window. Path efficiency, *Eff*, is quantified by the cumulative –log(LMI) of all edges along the path. The efficiency difference is calculated as $\Delta Eff = Eff_{\mathrm{Gln322-Leu546}} - Eff_{\mathrm{Gln322-Leu754}}$. "RepX-Y" means the Yth simulation window of replicate X. Each window spans 200 ns. Y starts from 11 because we focused on the data from 2 to 4 µs. Entries labeled with "NA" are cases where no path can be found or lack the data required to calculate $\Delta Eff$.

| | $Eff_{\mathrm{Gln322-Leu546}}$ | $Eff_{\mathrm{Gln322-Leu754}}$ | $\Delta Eff$ |
|---|---|---|---|
| Rep1-11 | 2.459 | NA | NA |
| Rep1-12 | 2.649 | 2.741 | -0.092 |
| Rep1-13 | 2.470 | 4.550 | -2.080 |
| Rep1-14 | 2.597 | 5.387 | -2.790 |
| Rep1-15 | 2.798 | 2.757 | 0.041 |
| Rep1-16 | NA | NA | NA |
| Rep1-17 | 2.911 | 3.185 | -0.274 |
| Rep1-18 | 2.595 | NA | NA |
| Rep1-19 | 2.978 | NA | NA |
| Rep1-20 | NA | 4.746 | NA |
| Rep2-11 | 3.263 | 1.844 | 1.418 |
| Rep2-12 | 2.854 | 1.930 | 0.924 |
| Rep2-13 | 2.961 | 5.465 | -2.504 |
| Rep2-14 | 3.332 | 7.403 | -4.071 |
| Rep2-15 | NA | NA | NA |
| Rep2-16 | NA | NA | NA |
| Rep2-17 | 3.351 | 6.164 | -2.813 |
| Rep2-18 | 2.046 | NA | NA |
| Rep2-19 | 1.676 | NA | NA |
| Rep2-20 | NA | NA | NA |
| Rep3-11 | NA | NA | NA |
| Rep3-12 | 2.262 | NA | NA |
| Rep3-13 | NA | NA | NA |
| Rep3-14 | 1.867 | NA | NA |
| Rep3-15 | 3.058 | 1.685 | 1.373 |
| Rep3-16 | 3.698 | NA | NA |
| Rep3-17 | NA | NA | NA |
| Rep3-18 | NA | NA | NA |
| Rep3-19 | 2.266 | NA | NA |
| Rep3-20 | 2.167 | NA | NA |

**Table S3.** Solvent-exposed residues' proxies related to recovering the experimental initiation loop and Leu546-directed cone. The residue entries are ordered in descending order of the corresponding root mean square fluctuation (RMSF) value. The RMSF calculation includes all catalytic domain residues: His147 to Ile839 (Figure S3). Shown here are only the results of the selected 73 residues. The residues highlighted in red are the top 20 with the highest Leu546 preference, i.e., the highest preference score calculated as the logarithm of the path occurrence ratios: $\log\left(\frac{Occr\_path_{546}}{Occr\_path_{754}}\right)$, whereas those in blue are the top 20 with the highest Leu754 preference, i.e., the lowest preference score. Note that in the cross-comparison, we excluded residues showing fewer than 1/6 of the maximum path occurrences (i.e., < 1000) towards both L546 and L754 to exclude inert ones in thermal energy transfer: Asp320 and Asn172. Residues selected by the cross-comparison (Figures 4b and S5a) are bold.

| Index | Residue index | RMSF | $Occr\_path_{546}$ | $Occr\_path_{754}$ | Occr_path ratio | Preference score |
|---|---|---|---|---|---|---|
| 1 | 320 | 3.497 | 900 | 400 | 2.250 | 0.352 |
| 2 | 455 | 3.086 | 2500 | 2500 | 1.000 | 0.000 |
| 3 | **322** | 2.240 | 2250 | 1400 | 1.607 | 0.206 |
| 4 | **318** | 2.213 | 2500 | 1200 | 2.083 | 0.319 |
| 5 | 174 | 2.039 | 100 | 100 | 1.000 | 0.000 |
| 6 | 277 | 2.003 | 3306 | 2600 | 1.272 | 0.104 |
| 7 | **454** | 1.977 | 3250 | 3900 | 0.833 | -0.079 |
| 8 | 596 | 1.938 | 0 | 0 | NA | NA |
| 9 | 391 | 1.910 | 1900 | 1800 | 1.056 | 0.023 |
| 10 | **278** | 1.882 | 3712 | 2100 | 1.768 | 0.247 |
| 11 | **439** | 1.750 | 2150 | 2600 | 0.827 | -0.083 |
| 12 | 440 | 1.578 | 100 | 0 | NA | NA |
| 13 | 393 | 1.540 | 4200 | 4400 | 0.955 | -0.020 |
| 14 | 172 | 1.510 | 200 | 300 | 0.667 | -0.176 |
| 15 | 606 | 1.437 | 500 | 500 | 1.000 | 0.000 |
| 16 | 210 | 1.410 | 0 | 0 | NA | NA |
| 17 | 814 | 1.383 | 2800 | 3000 | 0.933 | -0.030 |
| 18 | 594 | 1.362 | 4900 | 5200 | 0.942 | -0.026 |
| 19 | **363** | 1.362 | 1250 | 1500 | 0.833 | -0.079 |
| 20 | **438** | 1.318 | 3300 | 4400 | 0.750 | -0.125 |
| 21 | 394 | 1.285 | 4800 | 4800 | 1.000 | 0.000 |
| 22 | 236 | 1.275 | 0 | 200 | NA | NA |
| 23 | 298 | 1.270 | 3100 | 1100 | 2.818 | 0.450 |
| 24 | 378 | 1.250 | 300 | 500 | 0.600 | -0.222 |
| 25 | 389 | 1.222 | 0 | 0 | NA | NA |
| 26 | 211 | 1.198 | 0 | 0 | NA | NA |
| 27 | 718 | 1.166 | 4900 | 3400 | 1.441 | 0.159 |
| 28 | 364 | 1.147 | 0 | 100 | NA | NA |
| 29 | 176 | 1.143 | 1950 | 2300 | 0.848 | -0.072 |
| 30 | 593 | 1.091 | 4200 | 4100 | 1.024 | 0.010 |
| 31 | 471 | 1.075 | 1400 | 1200 | 1.167 | 0.067 |

| | | | | | | |
|---|---|---|---|---|---|---|
| 32 | 197 | 1.073 | 2700 | 5400 | 0.500 | -0.301 |
| 33 | 179 | 1.057 | 1350 | 1600 | 0.844 | -0.074 |
| 34 | 227 | 1.052 | 2950 | 4800 | 0.615 | -0.211 |
| 35 | 370 | 1.046 | 4650 | 5000 | 0.930 | -0.032 |
| 36 | 717 | 1.031 | 4700 | 3300 | 1.424 | 0.154 |
| 37 | 771 | 1.031 | 0 | 0 | NA | NA |
| 38 | 234 | 1.030 | 1636 | 4100 | 0.399 | -0.399 |
| 39 | 294 | 1.025 | 2850 | 1400 | 2.036 | 0.309 |
| 40 | 196 | 1.001 | 1200 | 5000 | 0.240 | -0.620 |
| 41 | 337 | 0.969 | 2450 | 1500 | 1.633 | 0.213 |
| 42 | 178 | 0.968 | 1800 | 2100 | 0.857 | -0.067 |
| 43 | 766 | 0.960 | 2500 | 2500 | 1.000 | 0.000 |
| 44 | 228 | 0.954 | 900 | 2400 | 0.375 | -0.426 |
| 45 | 602 | 0.935 | 5500 | 4700 | 1.170 | 0.068 |
| 46 | 284 | 0.903 | 4000 | 2600 | 1.538 | 0.187 |
| 47 | 336 | 0.892 | 3400 | 2800 | 1.214 | 0.084 |
| 48 | 424 | 0.885 | 0 | 0 | NA | NA |
| 49 | 334 | 0.883 | 2400 | 2300 | 1.043 | 0.018 |
| 50 | 180 | 0.872 | 500 | 100 | 5.000 | 0.699 |
| 51 | 425 | 0.864 | 750 | 400 | 1.875 | 0.273 |
| 52 | 326 | 0.832 | 4700 | 3000 | 1.567 | 0.195 |
| 53 | 230 | 0.820 | 1050 | 3600 | 0.292 | -0.535 |
| 54 | 380 | 0.817 | 5000 | 5500 | 0.909 | -0.041 |
| 55 | 368 | 0.795 | 2950 | 3300 | 0.894 | -0.049 |
| 56 | 760 | 0.787 | 300 | 200 | 1.500 | 0.176 |
| 57 | 774 | 0.782 | 2300 | 1400 | 1.643 | 0.216 |
| 58 | 759 | 0.766 | 3100 | 1800 | 1.722 | 0.236 |
| 59 | 825 | 0.766 | 1750 | 1200 | 1.458 | 0.164 |
| 60 | 329 | 0.763 | 1600 | 100 | 16.000 | 1.204 |
| 61 | 826 | 0.759 | 1900 | 700 | 2.714 | 0.434 |
| 62 | 233 | 0.733 | 1900 | 5100 | 0.373 | -0.429 |
| 63 | 670 | 0.732 | 1250 | 1800 | 0.694 | -0.158 |
| 64 | 331 | 0.714 | 2100 | 1900 | 1.105 | 0.043 |
| 65 | 282 | 0.700 | 3500 | 1892 | 1.850 | 0.267 |
| 66 | 668 | 0.676 | 2700 | 2300 | 1.174 | 0.070 |
| 67 | 662 | 0.670 | 0 | 0 | NA | NA |
| 68 | 427 | 0.666 | 2200 | 2000 | 1.100 | 0.041 |
| 69 | 663 | 0.666 | 0 | 0 | NA | NA |
| 70 | 182 | 0.664 | 3200 | 4200 | 0.762 | -0.118 |
| 71 | 382 | 0.641 | 3350 | 3300 | 1.015 | 0.007 |
| 72 | 823 | 0.600 | 2150 | 2900 | 0.741 | -0.130 |
| 73 | 831 | 0.553 | 2950 | 800 | 3.688 | 0.567 |
